# Differential and defective expression of Koala Retrovirus indicate complexity of host and virus evolution

**DOI:** 10.1101/211466

**Authors:** R.E Tarlinton, A.R. Legione, N. Sarker, J. Fabijan, J. Meers, L. McMichael, G. Simmons, H. Owen, J.M. Seddon, G. Dick, J.S. Ryder, F. Hemmatzedah, D.J. Trott, N. Speight, N. Holmes, M. Loose, R.D. Emes

## Abstract

Koala retrovirus (KoRV) is unique amongst endogenous (inherited) retroviruses in that its incorporation to the host genome is still active, providing an opportunity to study what drives this fundamental process in vertebrate genome evolution. Animals in the southern part of the natural range of koalas were previously thought to be either virus free or to have only exogenous variants of KoRV with low rates of KoRV induced disease. In contrast, animals in the northern part of their range universally have both endogenous and exogenous KoRV with very high rates of KoRV induced disease such as lymphoma. This paper uses a combination of sequencing technologies, Illumina RNA sequencing of “southern” (south Australian) and “northern” (SE QLD) koalas and CRISPR enrichment and nanopore sequencing of DNA of “southern” (South Australian and Victorian animals) to retrieve full length loci and intregration sites of KoRV variants. We demonstrate that koalas that tested negative to the KoRV *pol* gene qPCR, used to detect replication competent KoRV, are not in fact KoRV free but harbour defective, presumably endogenous, “RecKoRV” variants that are not fixed between animals. This indicates that these populations have historically been exposed to KoRV and raises questions as to whether these variants have arisen by chance or whether they provide a protective effect from the infectious forms of KoRV. This latter explanation would offer the intriguing prospect of being able to monitor and selectively breed for disease resistance to protect the wild koala population from KoRV induced disease.

## Introduction

Koalas (*Phascolarctos cinereus*) are an iconic marsupial species listed as vulnerable on the IUCN ‘red list’ of threatened species ^1^. While a large part of their ongoing population decline is due to habitat loss, two major disease threats, chlamydial infection and Koala Retrovirus (KoRV), are additionally limiting population viability ^2^. These infections are particularly prevalent in the northern regions of Australia, namely the states of Queensland and New South Wales, and less so in the south (South Australia, Victoria) ^3,4^.

Following European settlement, large koala populations across Australia declined significantly due to hunting in the 1890’s to 1920’s, with southern populations nearing extinction. During this time, small refuge populations were established on offshore Victorian islands and these koalas have been used subsequently to restock most of their former southern range. This southern population is genetically distinct from the northern animals ^5^ with a more limited genetic diversity^6^. The history of translocations in southern animals is complex but the original founder populations of French and Phillip Islands are thought to have been the source for most mainland Victorian animals with potential remnant populations of greater diversity in the Strzelecki ranges ^5^. The mainland Mount Lofty Ranges koala population in South Australia originates from koalas from both the Kangaroo Island population, populated by koalas from French Island ^7^ as well as koalas from Queensland and New South Wales ^5,8^.

Endogenous retroviruses (ERVs) are those that have become incorporated into their host’s genome. They are ubiquitous in vertebrate genomes and in some cases constitute up to 10% of total genome content ^9^. They are usually not functional viruses due to the accumulation of mutations and deletions but are often expressed at an RNA level, where they are thought to play a role in genomic regulation ^9 10,11^. They are known in some cases to provide essential functions to their hosts, such as the syncytin genes responsible for placental fusion in many species ^12,13^ as well as their role in stem cells, reproductive tissue and early embryos ^14^. However their effects on the host upon initial entry to the host genome are not clear. KoRV is part of a small group of unusual “modern” endogenous retroviruses (including Murine leukaemia virus, Feline leukaemia virus and Jaagsietke sheep retrovirus). These modern ERVs are replication competent and display considerable overlap with their exogenous infectious counterparts, including swapping of gene segments ^15,16^.

KoRV is one of the most recent entrants into any known mammalian genome, with estimates of integration time somewhere between 200 and 49,000 years ago ^17,18^.It is thought to have arisen from a recent species jump as its closest relatives are endogenous viruses in two subspecies of *Melomys burtoni* (the grassland mosaic tailed rat) in northern Australia and Indonesia ^19,20^, Gibbon ape leukaemia virus (GALV), a pathogenic exogenous virus that most likely arose as a spill over event from south east Asian rodents in the late 1960s ^21^ and Flying fox retrovirus, a very recently described exogenous virus of black flying foxes (*Pteropus alecto*) ^22 23^.

KoRV was originally identified during investigations into the high rates of lymphoid neoplasia (lymphoma and leukaemia) in Queensland koalas ^17^. Koalas with lymphoid neoplasia have significantly higher KoRV viral loads ^24 25^ and some strains of KoRV also influence the cytokine response profile of koala lymphocytes ^26^. Recent studies have indicated that somatic insertions of KoRV perturb oncogenes and underlie the very high rate of cancer in KoRV A positive animals ^27^. Multiple studies also indicate that high KoRV viral loads (in northern populations) or positive PCR status (in southern populations) ^28–32^ are linked to clinical chlamydial disease, probably as a factor of retroviral induced immunosuppression.

KoRV has been found in 100% of Queensland and New South Wales koalas but appears to have a lower prevalence in southern populations ^4,28,30,32–34^. The virus displays a high diversity in proviral copy number and integration sites between individuals and populations, with southern animals having lower copy numbers in their DNA ^24,35 4^. Somatic insertions are also apparent against a background of endogenous insertions in northern animals ^27^.

A number of sequence variants of the *env* gene region, which encodes the surface unit (SU) of the envelope protein (Env), have also been identified (Figure 1). These vary between individuals and resemble the viral quasispecies common to infectious retroviruses, with clades referred to as A to J ^34,36^. The originally identified virus is now known as KoRV A and appears to be present in all individuals that are KoRV-positive ^27,28,30,33,37^. Various koala genome sequencing studies indicate that only KoRV A is endogenised in northern animals with other variants present at lower than one copy/genome equivalent, indicating that they are not present in all tissues or cells of an animal ^27,35,38^. A recent study indicated that there may be one KoRV A locus shared amongst most (perhaps all) northern animals, which perhaps represents the original endogenisation event^27^. KoRV A infections in southern animals may represent genuine exogenous (infectious) virus as these are in many cases also present at less than one copy per genome equivalent ^4^. The non-A variants may also represent genuine exogenous (infectious) virus in both northern and southern animals, circulating independently with these present as low copy number/somatic insertions^27,38,39^, not detected in all animals ^29,30,34,40–42^ and display a pattern of detection in family groupings consistent with a maternally transmitted infection ^42 40 33 43^. Some caution is necessary in interpreting this however as phylogenetic analysis of the envelope variants from a variety of sequencing studies do not clearly indicate chains of transmission ^34,36,41^ and, by analogy with infectious retroviruses in other species (for instance FeLV in cats), many envelope sequence variants may arise from KoRV A within individual infections rather than transmitting from animal to animal ^15^ or may be transmitted as a co-infection with KoRV-A. This is particularly likely for many of the “D” group of variants that do not appear to be replication competent ^34,44,45^.

**Figure 1:**
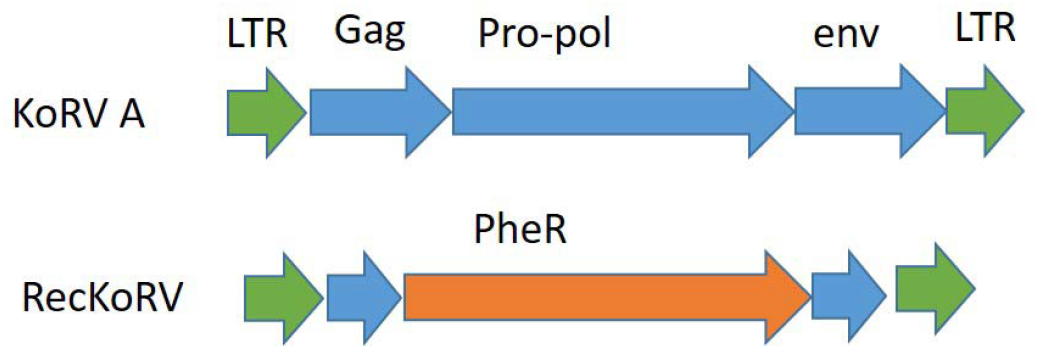
Cartoon of KoRVA and RecKoRV genetic sequence. KoRV LTRs are marked in green, KoRV sequences in blue, PhER sequences in orange.

**Figure 2:**
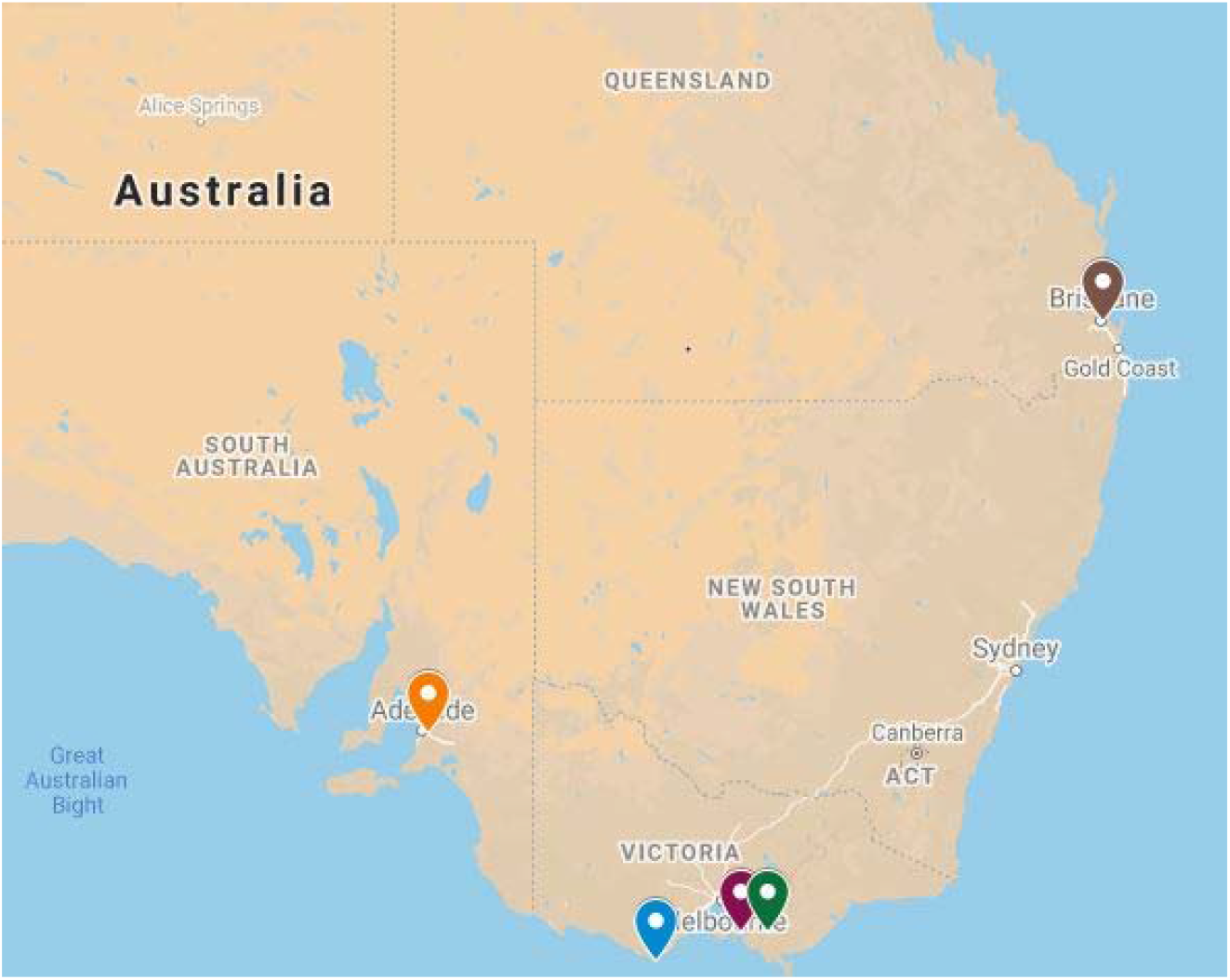
Map of the locations of the animals sampled in this study. Mt Lofty Ranges orange drop (SA), Cape Otway blue drop, French Island purple drop, Strezlecki ranges green drop (VIC), SE QLD brown drop (QLD) (map created with Google maps)

There has been much debate as to whether the B/J variant, which displays a different receptor usage to KoRV A is more pathogenic as these variants have been epidemiologically linked with clinical disease in some studies but not others ^46 47 29 33^. This may however be a factor of the sensitivity of diagnostic methods used as at least one study has demonstrated that koalas with higher viral loads display greater quasispecies diversity and are more likely to test positive on PCR based tests for non-A variants ^36^. That study also demonstrated that viral diversity is much higher in RNA (actively replicating virus) than DNA (copies inserted either endogenously or from initial infection) from the same animal.

Genomic sequencing studies have also demonstrated that there are a number of other older endogenous retroviruses and transposable elements within the koala genome ^6,38,48,49^. One of these, phascolarctid endogenous retroelement (PhER), is found frequently in northern koala genomes in recombination with KoRV. These recombinant KoRV “RecKoRV” structures typically consist of the 5’ LTR and 5’ end of the KoRV *gag* gene, approximately 5 Kb of the 3’ end of PhER and its LTR, followed by the 3’ end of the KoRV env gene and KoRV 3’LTR ^38,49^ (Figure 1). There appear to be multiple variants of these that arise from very similar recombinations at particular points in the KoRV/PhER genomes. They are not shared between all animals but do display some geographical clustering in loci that are shared between individuals and may be absent in some populations ^49^. Variants of KoRV A with large indels or “Solo LTRs” (where the middle part of the virus is spliced out during cellular DNA replication) are also seen ^38^.

This study reports the presence of RecKoRV variants in Southern animals that do not carry KoRV A. These variants appear to be a different genetic lineage to that present in northern animals and to be present (though not fixed) in all animals tested from multiple Victorian and South Australian populations, including the founder population on French Island.

## Methods

### Ethics

Ethical approval for this study was granted by the University of Queensland Animal Ethics Committee, permit number ANFRA/SVS/461/12, the Queensland Government Department of Environment and Heritage Protection permit number WISP11989112, the University of Adelaide Animal Ethics Committee permit number S-2013-198 and the South Australian Government Department of Environment, Water and Natural Resources Scientific Research Permit Y26054, the University of Nottingham School of Veterinary Medicine and Science Clinical Ethics Research Panel, and Department of Environment and Primary Industries (Victoria, Australia) (Research Permit 10006924).

### Samples for DNA sequencing

DNA sequencing was performed on samples from five southern koalas (Table 1 B). Spleen samples from three wild Victorian animals were collected at necropsy as outlined in ^30^. Liver samples were collected from one 3 year old female South Australian koala housed in a zoological park in the UK that had been recently imported from an Australian captive population derived from the Mt Lofty ranges and Kangaroo Island population in SA. This animal died of the kidney disease oxalate neprosis with samples of liver collected at post mortem and stored at −20°C until DNA extraction and sequencing. Lymph node samples were collected from one wild Mt Lofty (SA) that died as a result of dog attack as described in ^25^.

**Table 1:**
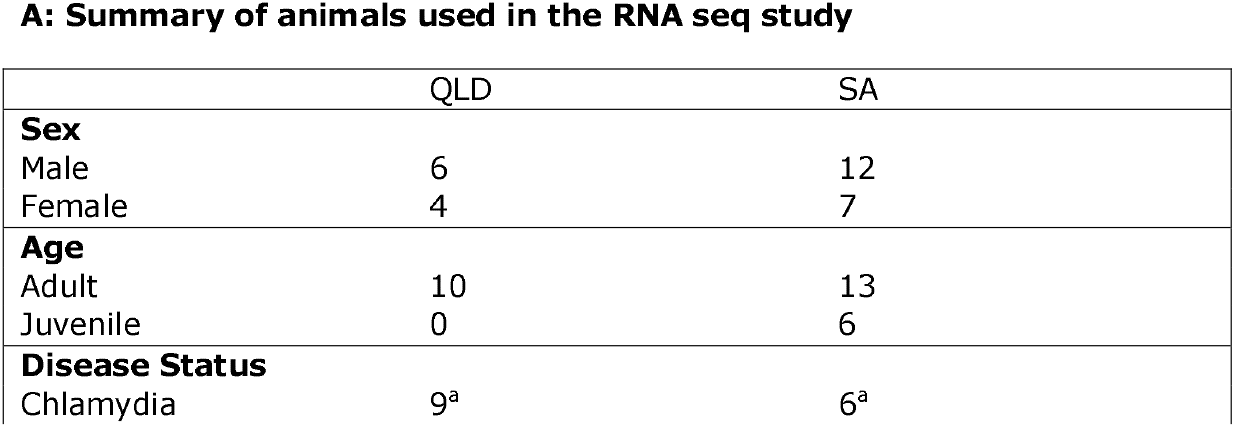
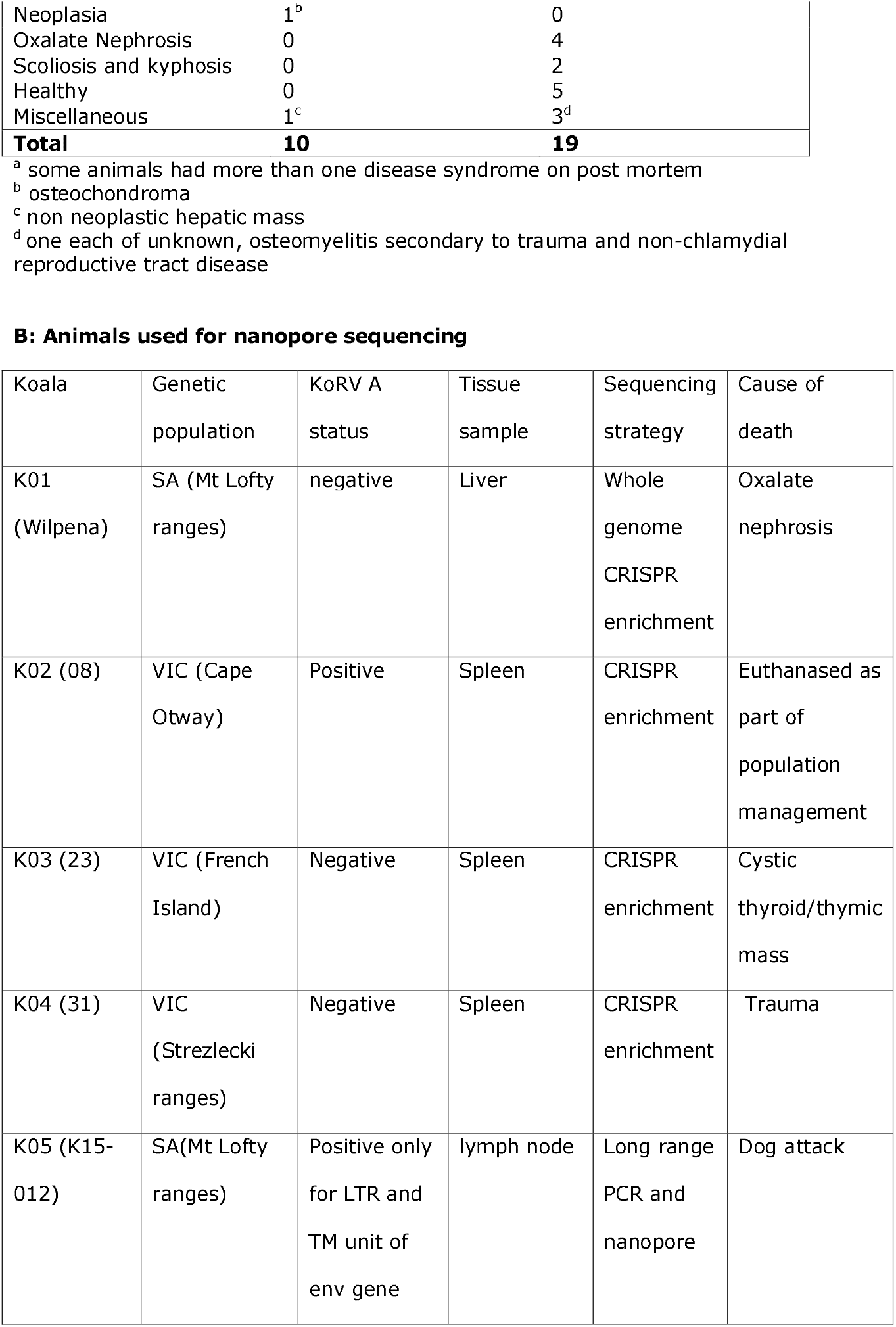
Details of the koalas used in this study.

**A: Summary of animals used in the RNA seq study**
|  | QLD | SA |
| --- | --- | --- |
| **Sex** |  |  |
| Male | 6 | 12 |
| Female | 4 | 7 |
| **Age** |  |  |
| Adult | 10 | 13 |
| Juvenile | 0 | 6 |
| **Disease Status** |  |  |
| Chlamydia | 9<sup>a</sup> | 6<sup>a</sup> |
| Neoplasia | 1<sup>b</sup> | 0 |
| Oxalate Nephrosis | 0 | 4 |
| Scoliosis and kyphosis | 0 | 2 |
| Healthy | 0 | 5 |
| Miscellaneous | 1<sup>c</sup> | 3<sup>d</sup> |
| **Total** | **10** | **19** |

<sup>a</sup> some animals had more than one disease syndrome on post mortem
<sup>b</sup> osteochondroma
<sup>c</sup> non neoplastic hepatic mass
<sup>d</sup> one each of unknown, osteomyelitis secondary to trauma and non-chlamydial reproductive tract disease

B: Animals used for nanopore sequencing
| Koala | Genetic population | KoRV A status | Tissue sample | Sequencing strategy | Cause of death |
| --- | --- | --- | --- | --- | --- |
| K01 (Wilpena) | SA (Mt Lofty ranges) | negative | Liver | Whole genome CRISPR enrichment | Oxalate nephrosis |
| K02 (08) | VIC (Cape Otway) | Positive | Spleen | CRISPR enrichment | Euthanased as part of population management |
| K03 (23) | VIC (French Island) | Negative | Spleen | CRISPR enrichment | Cystic thyroid/thymic mass |
| K04 (31) | VIC (Strezlecki ranges) | Negative | Spleen | CRISPR enrichment | Trauma |
| K05 (K15-012) | SA(Mt Lofty ranges) | Positive only for LTR and TM unit of env gene | lymph node | Long range PCR and nanopore | Dog attack |

### Samples for RNA seq

Samples were collected from wild-rescued koalas euthanised for clinical reasons and submitted for post-mortem examinations from South East Queensland (Greater Brisbane) (n=10) and South Australia (Mount Lofty Ranges) (n=19). Age was determined by dentition and the amount of wear on the upper premolar ^50^ (Table 1 A). Full details of these animals are presented in ^6^. Submandibular lymph nodes were collected within 2-6 hours of death into RNALater^®^ and stored at −80°C. Where possible, blood was collected into EDTA prior to euthanasia (BD vacutainer) with whole blood and plasma added to RNA later as per previous studies ^51^ kept at −80°C. Of the ten koalas from South East Queensland (QLD), six were male and four female and all were adults, with a tooth wear class (TWC) 4 or 5. Nineteen koalas were sampled from the Mount Lofty Ranges, South Australia (SA); seven female and 12 male. Six were juvenile (TWC 1 or 2) and 13 were adults (TWC 3 or 4).

### Nanopore Sequenced animals

#### DNA extraction for nanopore sequencing

Genomic DNA was extracted from frozen liver/spleen tissue that had been ground into a fine powder under liquid nitrogen. The Qiagen Genomic Tip (100/G) kit (Qiagen; 10243) was used to extract DNA from 100 mg of tissue powder. DNA was quantified using the Qubit Fluorometer (Thermo Fisher Scientific) and the Qubit dsDNA BR Assay Kit (Thermo Fisher Scientific; Q32853) and the molecular weight was assessed using the Agilent TapeStation 4200 and the Agilent Genomic DNA ScreenTape Assay (Agilent; 5067-5365 and 5067-5366). A sequencing library was prepared using the Genomic DNA by Ligation Kit (Oxford Nanopore Technologies; SQK-LSK109) and run on a PromethION flow cell (Oxford Nanopore Technologies; FLO-PRO002) for 72 hours on a PromethION beta sequencer (Oxford Nanopore Technologies).

#### Nanopore Sequencing for KoRV insertions

Cas9-mediated PCR-free enrichment was performed to identify individual KoRV insertion sites. Genomic DNA was also extracted as described above or was extracted from spleen tissue, that had been stored in RNAlater (ThermoFisher) at −80°C, using the Qiagen PureGene DNA extraction Kit (Qiagen; 158445).

Genomic DNA was dephosphorylated to inhibit binding of Oxford Nanopore sequencing adapters to non-specific DNA fragments. Six custom Alt-R CRISPR-Cas9 crRNA (Integrated DNA Technologies) were used to form Cas9 ribonucleoprotein complexes (RNPs) that would facilitate strand-specific cleavage at target sites within KoRV. Cleaved ends were simultaneously dA-tailed to facilitate directional ligation of sequencing adapters and enrich for reads initiating at these crRNA cleavage sites. Lyophilized crRNA were reconstituted to 100 uM TE (pH7.5) and pooled in equimolar amounts. Cas9-mediated enrichment, sequencing library preparation and sequencing were then performed according Oxford Nanopore Technologies Cas-mediated PCR-free enrichment protocol (Version: ENR_9044_v1_xxxx_08Aug18); and each library was run on a separate MinION flow cell (Oxford Nanopore Technologies; FLO-MIN106 R9.4.1) on the GridION X5 Mk1.

#### Nanopore Sequencing of PCR amplicons

PCR amplification was conducted using the primer set KRV R2 forward (ATCTACCCGGAGACGGACAG) and reverse (GCCGGTACCTATACCTGCTG) ^25^ to amplify an approximately 6kb fragment of the KoRV genome from extracted genomic DNA from the SA koala K15-012. A sequencing library was prepared using the Rapid Sequencing Kit (Oxford Nanopore Technologies; SQK-RAD004) and run on a MinION flow cell (Oxford Nanopore Technologies; FLO-MIN106D) for 36 hours on a MinION sequencer (Oxford Nanopore Technologies).

#### Sequence Assembly and Mapping

Nanopore sequences were basecalled using guppy and reads that passed the default read filtering metrics were obtained. Reads for each koala were mapped to the KoRV retrovirus reference genome (Genbank Accession number: AF151794) using minimap2 ^52^ and samtools ^53^. Read mapping was visualised using Geneious Prime software (Biomatters, New Zealand) and reads were truncated to retain regions upstream and downstream of the KoRV genome. These truncated reads were then mapped against the koala reference genome assembly (Genbank Accession number: GCA_002099425.1) using minimap2 with no secondary hits allowed. The mapped reads were visualised in Genious Prime to identify the directionality of the insert, whether the insert potentially interrupted coding regions of the koala genome, and identify upstream genes that could be influenced by insertion. Additionally, reads were mapped to a sequence of PhER ^54^.

All reads mapping to KoRV for each koala were assembled using flye ^55^ in order to obtain a consensus insert assembly. Additionally, reads that mapped to individual contigs of the koala reference genome, representing individual insert sites, were extracted and assembly was also attempted using flye.

### Animals for RNAseq

#### RNA preparation for RNAseq

Total RNA was extracted from lymph nodes using an RNeasy Mini kit with on column DNAase1 digestion (Qiagen). RNA quantity and quality were assessed via anXpose spectrophotometer (Bioke) and Agilent 2100 Bioanalyzer. mRNA was prepared for sequencing using the Illumina TruSeq stranded mRNA library prep kit and 100 base pair, paired end sequencing was performed on an Illumina HiSeq. Details of the koalas, sample quality and read quantity are provided in Supplementary data 1.

#### RNA and DNA extraction for qPCR/PCR (SA and QLD animals)

DNA was extracted from 100 μL of EDTA blood using a DNeasy blood and tissue kit (Qiagen). Where available RNA was extracted from plasma Using the QIAMp Viral RNA mini kit with on-column Qiagen RNase free DNAse digestion. The extracted RNA and DNA was stored at −80°C for RT-PCR (RNA) and PCR (DNA) as required.

#### KoRV qPCR

The presence of KoRV provirus for individual gene segments was assessed by qPCR for the KoRV A *pol* gene (the standard KoRV diagnostic assay)^31^ on DNA extracted from whole blood as reported in ^25^.

#### KoRV genome coverage

To reduce mis-mapping due to the abundance of highly repetitive long terminal repeat sequences, the adapter-trimmed fastq files were first mapped using Hisat2 ^56^ to the isolated Long Terminal Repeat (LTR) region of the koala KoRV type sequence (accession AF151794). LTR depleted reads were then mapped to representative sequences of KoRV A and RecKoRV derived from the koala reference genome (KoRV45 and RecKoRV6 Supplementary data 2 and 3) ^56^. Per-base coverage was determined from bam files for each isolate using samtools version 1.3.1 depth (with parameters –aa –q 10 –d 20000).

#### KoRV envelope variant gene expression

To quantitate the expression of KoRV envelope variants, LTR depleted reads for individual koalas were pseudoaligned to the *gag, pol* and *env* genes of KoRVA (accessions AAF15097.1_1, AAF15097.1_2 and AAF15097.1_3 respectively) and the first 575 nucleotides of the envelope variants of the non-A KoRV variants B-I (accessions AB822553.1, AB828005.1, AB828004.1, KX588043.1, KX587994.1, KX587961.1, KX588036.1 and KX588021.1 respectively) and the 3’ overlap of PhER/KoRV in RecKoRV using Kallisto ^57^. These nucleotides correspond to the hypervariable region of the *env* gene that is used in KoRV envelope variant classification.

#### Data Availability

KoRV sequence data (as fasta formatted data) are available from adac figshare [https://figshare.com/authors/Adac_uon_Adac_uon/566308]. Raw RNA sequence reads available in FASTQ format at ENA with the accession number PRJEB21505. Nanopore sequence data is available via accession number PRJNA770362.

## Results

RNA from submandibular lymph nodes from 10 QLD and 19 SA animals was subjected to paired end illumina sequencing (HiSeq 100bp) and was mapped to representative KoRVA and RecKoRV sequences from the koala reference genome (Figure 3). Demographic data for individual animals are presented in Supplementary data set 1.

**Figure 3:**
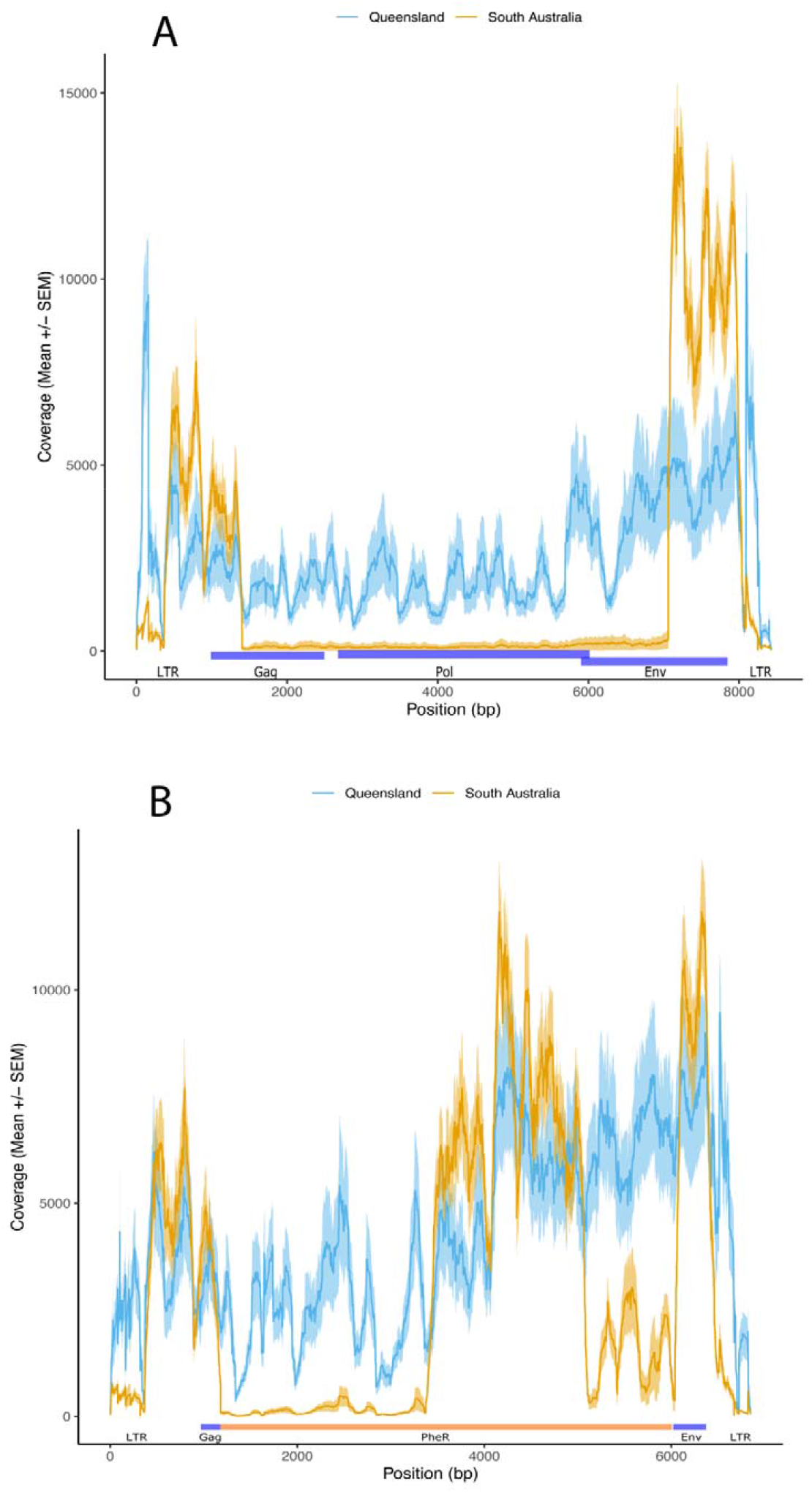
**Panel A:** Coverage of reads mapped to a representative sequence of KoRV A from the koala reference genome. For each group the mean normalised coverage ((per position coverage/total coverage) x 1×10^6^) is represented by a line and +/- the standard error is shaded around the mean. QLD samples in blue, SA in orange. KoRV genomic regions are marked underneath the read maps with blue bars, these regions are: 5’ LTR, gag, pol, env, LTR for KoRV A. **Panel B:** Coverage of reads mapped to a representative sequence of RecKoRV from the koala reference genome. For each group the mean normalised coverage ((per position coverage/total coverage) x 1×10^6^) is represented by a line and +/- the standard error is shaded around the mean. QLD samples in blue, SA in orange. RecKoRV genomic regions are marked underneath the read maps with blue bars, these regions are: 5’ LTR, gag portion, PhER, env portion, 3’ LTR.

When normalised for total mapped read depth, coverage was very similar for both the SA and QLD groups of koalas across the ends of the KoRV genomes (LTR-gag, and env-LTR). However, between positions 1389 and 7124 of the KoRV A sequence the SA group showed a mean coverage of < 10% of the QLD group suggesting that part of gag, all of *pro-pol* and part of the *env* genes were largely missing in the RNA transcripts, with six SA koalas not expressing this region at all (Figure 3 A). The target site of the standard KoRV *pol* qPCR used in most studies is contained within region ^51^. Data from other publications from this sample cohort indicate that some of SA animals were KoRV PCR positive for the proviral *pol* gene (and other genes) suggesting that at least partial proviruses for this region were present but were expressed at levels undetectable in the transcriptome^25^.

The higher number of reads in the *env* and LTR regions of the QLD animals can be explained by the presence of spliced *env* transcripts in addition to full length genomic transcripts as has been reported by other groups ^54^, although these are not detected as complete individual transcripts by the mapping methods used in this study.

Mapping of the reads to RecKoRV demonstrated relatively even coverage from the QLD animals, However there was little to no coverage of the 3’ portion of the PhER segment of RecKoRV in the SA animals, indicating that while there are RecKoRV sequences in the SA animals these likely differ in sequence from those in the genome animal (Figure 3 B)

Pseudomapping of the sequence reads to the KoRV A genome (complete *gag, pro-pol* and *env* genes) and type sequences of the hypervariable region of the *env* gene (base pairs 6000-6575 of KoRV A) of each of the previously identified KoRV envelope variants (KoRV A to I as per the classification scheme used in Chappell *et.al*. 2016 ^34^) demonstrated that while QLD koalas had multiple envelope variants within individuals, SA animals had far lower KoRV envelope variant diversity. Significantly higher expression was observed for KoRV A,B,D,E and G variants in QLD compared to SA samples (unpaired t-test with unequal variance) (Figure 4 and Supplementary data 4). It was observed that QLD animals were older (mean tooth wear class 4.22 95% CI 3.88-4.56) than SA (mean tooth wear class 3.05 95% CI 2.58-3.52) and so age may confound KoRV expression comparisons. When the same test was repeated for samples from koalas with the same tooth class 4 (7 QLD 8 SA samples), expression of A,B,E and G variants remained significantly different between locations (Supplementary data 5), supporting the finding that KoRV env expression is significantly higher in the QLD than the SA populations. Eleven out of nineteen SA animals (58%) had KoRV A. Six of these koalas had only KoRV A reads (Figure 4 and Table 2). Four animals had reads for KoRV A and one other variant only (D or E). Two animals had reads for KoRV E but no detectable reads for any other variant (including KoRV A). Only one SA koala (Z Table 2) had counts comparable to the QLD cohort with a similar range of variants (A, B, C, D, E, F, G, I), while the rest had counts that were <10% of the QLD koalas. *Pol* gene counts were also similarly considerably lower in the SA koalas than the QLD group. Relative expression as estimated count values for individual animals for each gene region and KoRV envelope variant are presented in Supplementary data 4.

**Figure 4:**
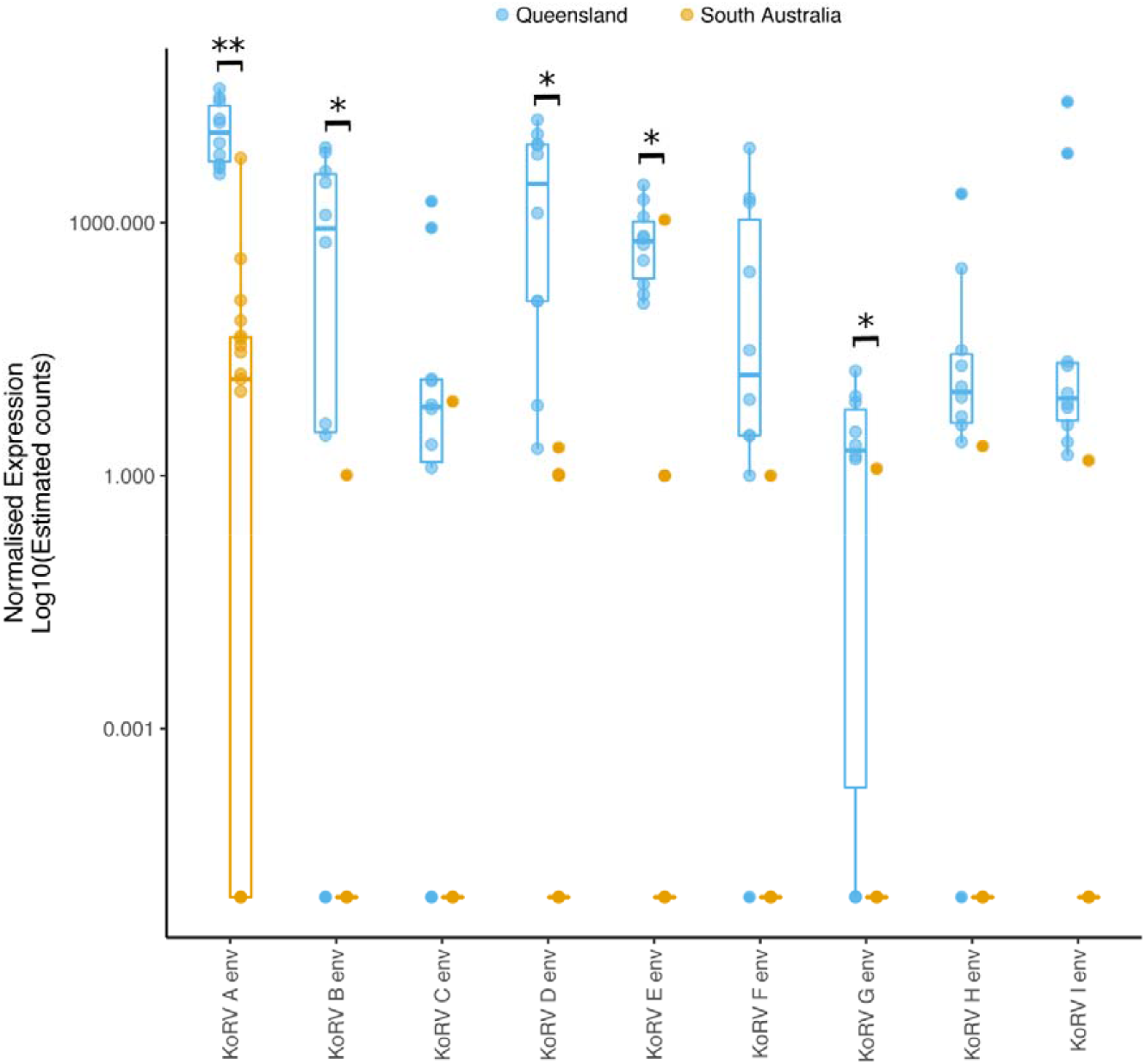
Normalised expression Log10(estimated counts) of KoRV A complete *env* gene and the 575 nucleotides of the hypervariable region of the envelope variants (B-I). Box and whisker plots show the median and interquartile ranges (box) and minimum/maximum expression (whiskers) of groups. Data for individual animals within a group are shown by circles. QLD animals in blue and SA animals in orange. *Env* variants with significantly different expression between QLD and SA groups marked with black bars (** = P<0.001, *= P<0.005)

**Table 2:** KoRV variant expression of individual animals.

| ID | Location<sup>a</sup> | Sex | Age<sup>b</sup> | Provirus PCR | KoRV variants<sup>c</sup> |
| --- | --- | --- | --- | --- | --- |
| A | QLD | M | 4 | + | ALL |
| B | QLD | M | 4 | + | ALL |
| C | QLD | F | 4 | + | ALL |
| D | QLD | F | 4 | + | ALL |
| E | QLD | F | >3 | + | A, C, D, E, F, G, H, I |
| F | QLD | M | 4 | + | A, D, E, I |
| G | QLD | M | 5 | + | A, B, C, D, E, F, H, I |
| H | QLD | M | 4 | + | A, B, D, E, F, G, H, I |
| I | QLD | F | 4 | + | ALL |
| J | QLD | M | 5 | + | A, B, C, D, E, F, H, I |
| K | SA | F | 4 | + | A |
| L | SA | M | 3 | + | A |
| M | SA | M | 2 | + | N<sup>d</sup> |
| N | SA | M | 4 | - | N |
| O | SA | M | 3 | + | N |
| P | SA | F | 3 | + | N |
| Q | SA | M | 4 | + | A |
| R | SA | M | 2 | + | A |
| S | SA | M | 2 | + | N |
| T | SA | M | 2 | + | A, D |
| U | SA | F | 4 | + | E |
| V | SA | M | 4 | + | E |
| W | SA | F | 4 | + | N |
| X | SA | F | 3 | - | A, D |
| Y | SA | F | 4 | - | A |
| Z | SA | M | 3 | + | A, B, C, D, E, F, G, I |
| A1 | SA | F | 1 | + | A, E |
| A2 | SA | M | 2 | - | A, D |
| A3 | SA | M | 4 | - | A |

<sup>a</sup> Population location: QLD – Queensland; SA – South Australia
<sup>b</sup> Age determined by dentition and the degree of wear of the upper pre-molar (Martin *et al.* 1999)
<sup>c</sup> KoRV variants determined by KoRV transcripts; ALL = all published variants (A to I)
<sup>d</sup> N= no env hypervariable region detected.

Mapping of CRISPR enriched nanopore sequences from koala samples to the KoRV reference genome identified a clear drop in coverage across the main portion of the genome. This went from base 450 in the *gag* coding region to base 1134 in the *env* coding region, or bases 1411 - 7040 across the KoRV-A reference genome (Figure 5). Mapping to a PhER assembly identified improved coverage, but there were still clear regions of near zero coverage in the mid-region of the reference (Figure 5). Importantly the three samples that had previously tested negative to KoRV using conventional PCR targeting the *pol* gene (Koalas 01 03 and 04) both had reads mapping to KoRV, but no coverage in the region of the PCR targets. Alignment of PhER and KoRV A from the (northern) reference genome animal and the sequence variants found in the southern animals is presented in Supplementary data 6. Assembly of reads that mapped to the koala reference genome generated 17 contigs containing RecKoRV variants (8 from K01-SA1-CRISPR, 7 from K01-SA1-WG, and 1 each from K03-Vic23 and K04-Vic31). The general structure of these inserts were similar across the assemblies besides a ~579 bp gap at the 5’ end at the interruption of the KoRV gag gene. Aligning all read sets back to one of the RecKoRV variants from Koala 01 showed that this insert was present across all koala samples (Figure 5).

**Figure 5.**
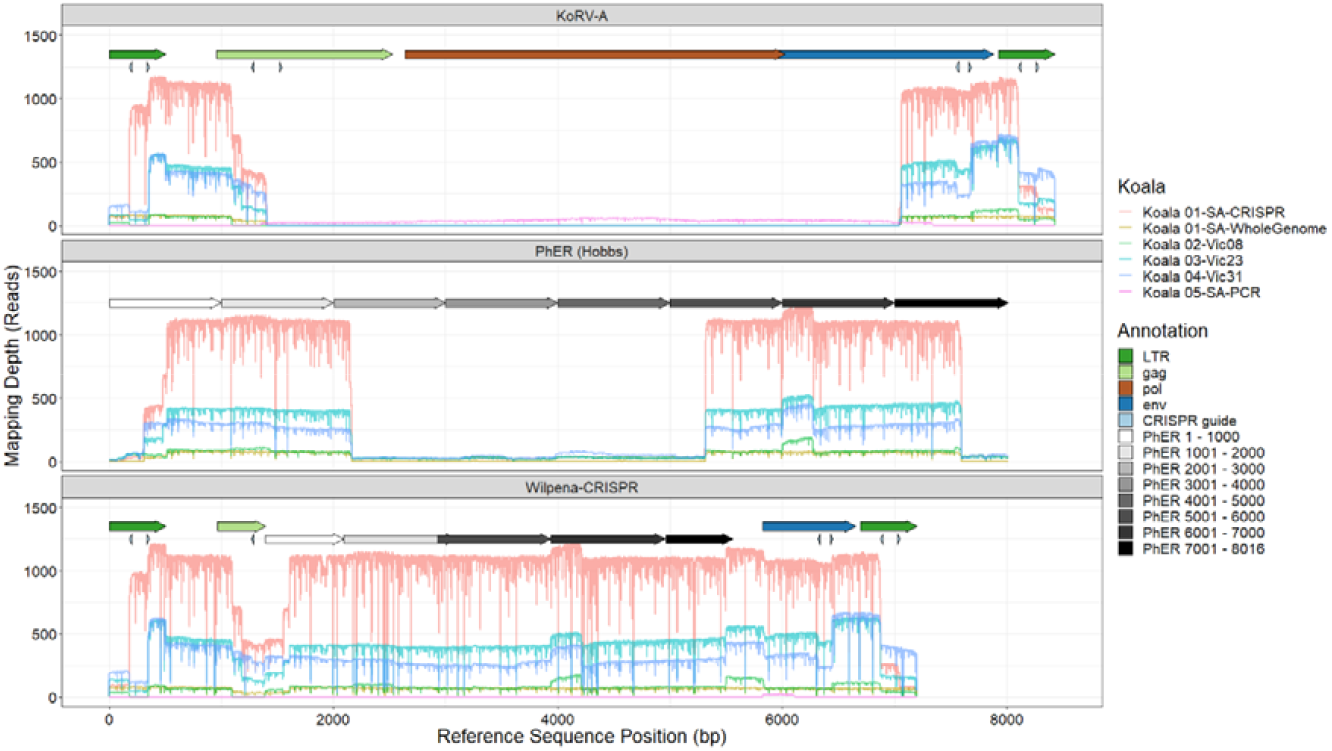
Coverage maps of Nanopore reads mapped to three different reference sequences (KoRV-A, PhER (Hobbs), Wilpena-CRISPR) using minimap2. Annotation arrows represent locations of coding domains from KoRV-A (in colour), CRISPR guide oligos, and genome regions of PhER in 1000 bp increments (greyscale) to highlight the insertion within the recKoRV assembly Wilpena-CRISPR. Note that the genomes do not align in the figure and base positions are relative to the reference genome in each plot.

Mapping of CRISPR enriched nanopore sequences from four koala samples identified potential KoRV insert locations on 30 koala reference genome contigs (filtering this to require at least five reads mapping at the same site in at least one koala to constitute an insertion point). The data from koala 5 could not be mapped in this way as the PCR and sequencing strategy excluded the insertion sites. A summary of insert sites and read mapping is available in Table 3. Of the predicted insertion points (Figure 6), eight were shared between samples, with koala 1 sharing insert sites with koalas 3 (2 contigs) and 4 (1 contig), and koala 3 sharing sites with koala 2 (1 contig) and 4 (4 contigs). No insertion sites were shared between all koalas

**Figure 6:**
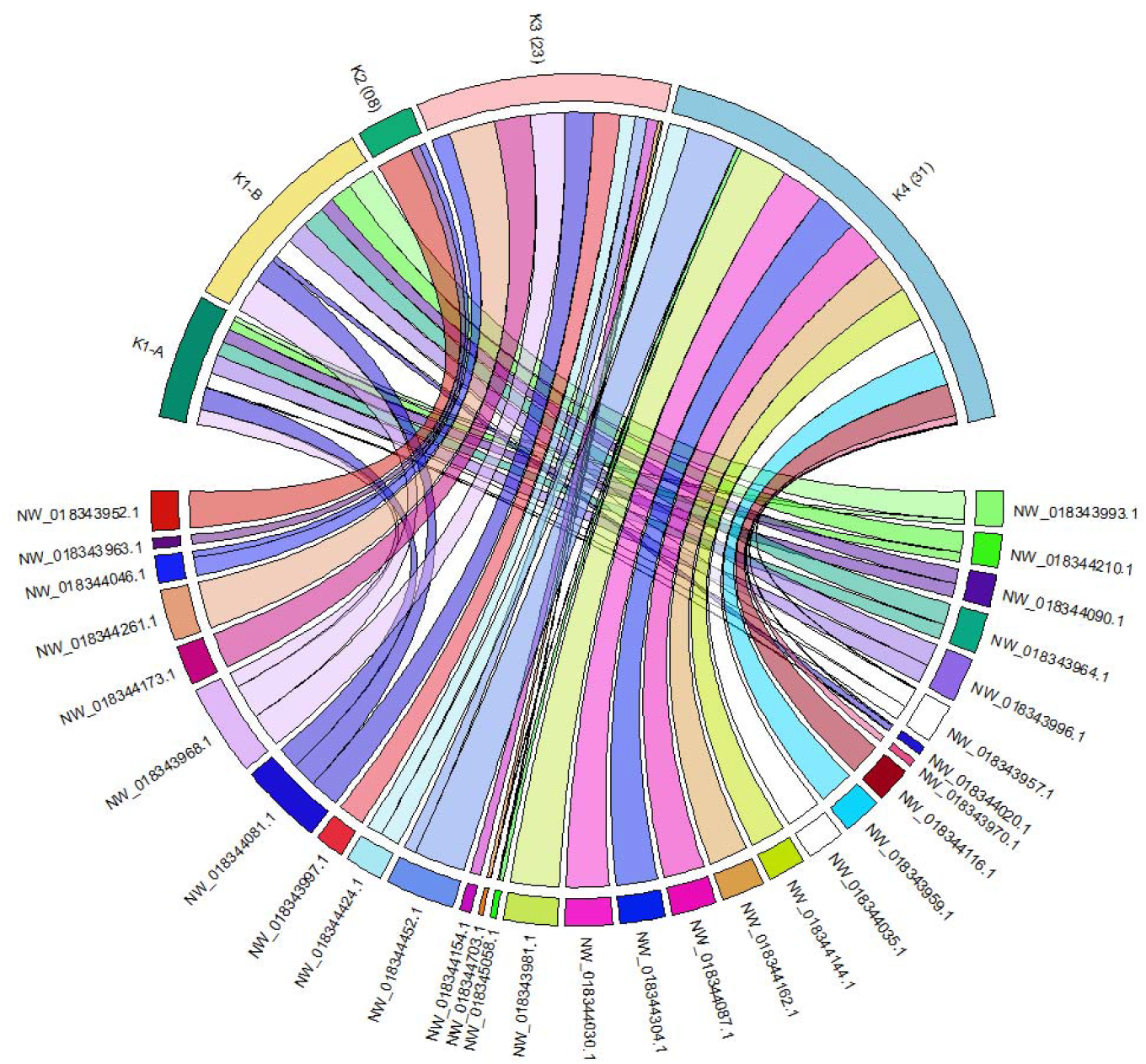
Circos plot of the number of reads mapping to koala reference genome contigs, highlighting the shared insert points between koalas 1 – 4.

**Table 3.** Summary information of total Nanopore reads matching to the koala reference genome

| Sample | Reads mapped to KoRV | Reads mapped to koala genome | Insertion Sites | Median (range) reads mapped per site |
| --- | --- | --- | --- | --- |
| Koala 1 – whole genome | 156 | 152 | 14 | 10 (1 – 26) |
| Koala 1 – CRISPR enrichment | 2488 | 272 | 16 | 14.5 (1 – 47) |
| Koala 2 | 275 | 72 | 3 | 13 (11 – 48) |
| Koala 3 | 1512 | 323 | 18 | 10 (1 – 63) |
| Koala 4 | 1699 | 609 | 25 | 5 (1 – 70) |
| Koala 5 | 156 | NA | NA | NA |
| Total | 6286 | 1428 | 56 | 8 (1 – 116) |

NA – Koala 5 nanopore reads generated using long-range PCR of KoRV primers, and did
not overlap the koala genome

An outline of interrupted genes, or genes downstream of KoRV insert sites, is presented in Table 4. Of the 30 insertion sites determined by mapping reads to the koala reference genome, 10 occurred within annotated genes, typically in predicted introns.

**Table 4.** Summary of insertion sites in the koala reference genome (Genbank Accession number: GCA_002099425.1) identified by mapping reads with minimap2

| Contig | Reads mapped | K01-WG | K01-CR | K2 (08) | K3 (23) | K4 (31) | Comment on Insert Site |
| --- | --- | --- | --- | --- | --- | --- | --- |
| NW_018343952.1 | 48 | 0 | 0 | 48 | 0 | 0 | Insert within MAP2K5 gene |
| NW_018343957.1 | 49 | 20 | 28 | 0 | 0 | 1 | Insert at different locations on contig between K1 and K4; K1 insert ~86 kb upstream of XR_002328485.1 Lnc RNA (potential interaction with VCAN gene) |
| NW_018343959.1 | 47 | 0 | 0 | 0 | 0 | 47 | Insert within MPP4 gene |
| NW_018343963.1 | 13 | 0 | 0 | 13 | 0 | 0 |  |
| NW_018343964.1 | 50 | 20 | 30 | 0 | 0 | 0 | ~36 kb upstream of RAB3GAP2 gene |
| NW_018343968.1 | 116 | 21 | 47 | 0 | 48 | 0 | ~150 kb upstream of STX6 gene |
| NW_018343970.1 | 5 | 0 | 0 | 0 | 0 | 5 | ~60 kb upstream of LOC110207063 gene |
| NW_018343981.1 | 68 | 0 | 0 | 0 | 0 | 68 | Insert within LOC110209428 gene |
| NW_018343993.1 | 45 | 7 | 38 | 0 | 0 | 0 | ~18 kb upstream of LOC110211657 gene |
| NW_018343996.1 | 56 | 26 | 30 | 0 | 0 | 0 | ~7 kb upstream of LOC110212362 gene |
| NW_018343997.1 | 32 | 0 | 0 | 0 | 32 | 0 | ~113 kb upstream of COMMD6 |
| NW_018344020.1 | 11 | 1 | 8 | 0 | 0 | 2 | ~789 kb gap between mapping of upstream and downstream regions |
| NW_018344030.1 | 58 | 0 | 0 | 0 | 0 | 58 | Insert within LOC110217408 gene |
| NW_018344035.1 | 54 | 0 | 0 | 0 | 0 | 54 | Insert within LOC110217834 LncRNA |
| NW_018344046.1 | 32 | 0 | 0 | 11 | 21 | 0 | Insert in CPA6 gene |
| NW_018344081.1 | 91 | 20 | 34 | 0 | 37 | 0 | Insert within TSPAN5 gene |
| NW_018344087.1 | 54 | 0 | 0 | 0 | 0 | 54 | ~18 kb upstream of BLOC1S6 gene |
| NW_018344090.1 | 40 | 19 | 21 | 0 | 0 | 0 |  |
| NW_018344116.1 | 45 | 0 | 0 | 0 | 0 | 45 | ~1 kb upstream of LOC110193889 LncRNA |
| NW_018344144.1 | 49 | 0 | 0 | 0 | 0 | 49 | ~80 kb upstream of HOOK3 |
| NW_018344154.1 | 15 | 0 | 0 | 0 | 15 | 0 | Insert within PITPNM2 gene |
| NW_018344162.1 | 53 | 0 | 0 | 0 | 0 | 53 | ~60 kb upstream of TM4SF20 gene |
| NW_018344173.1 | 47 | 0 | 0 | 0 | 47 | 0 | ~17, 20, 21, & 22 kb, upstream of tRNA-GCC, tRNA-GUC, tRNA-CUC, and LOC110197942 gene (predicted to encode heat shock 70 kDa protein 6-like) respectively |
| NW_018344210.1 | 42 | 13 | 29 | 0 | 0 | 0 | ~134 kb upstream of CXXC4 gene |
| NW_018344261.1 | 63 | 0 | 0 | 0 | 63 | 0 | Insert in PPFIBP1 gene |
| NW_018344304.1 | 58 | 0 | 0 | 0 | 0 | 58 | Insert within DCLK1 gene |
| NW_018344424.1 | 50 | 0 | 0 | 0 | 21 | 29 | Insert sites ~7 kb apart in different koalas |
| NW_018344452.1 | 88 | 0 | 0 | 0 | 18 | 70 | ~18 kb gap between mapping of upstream and downstream regions |
| NW_018344703.1 | 6 | 0 | 0 | 0 | 5 | 1 | Small genome contig (~41 kb) |
| NW_018345058.1 | 10 | 0 | 0 | 0 | 4 | 6 | Small genome contig (~31 kb) |
| NW_018345540.1 | 5 | 0 | 0 | 0 | 5 | 0 | Small genome contig (~14 kb) |
| **Totals** | 1400 | 147 | 265 | 72 | 316 | 600 |  |

## Discussion

The findings of the current study suggest that KoRV infection involves a more complex host-viral relationship than previously recognised, particularly in SA and Victorian koalas. Other studies have shown differences between northern and southern koala populations in the prevalence of KoRV infection, levels of KoRV proviral and viral loads and disease burden^25,58^. This study has revealed additional viral factors that indicate these population differences are more complicated than merely presence or absence of virus and virus load.

The results of this study were unexpected. Instead of these southern animals having demonstrably no KoRV as expected from a preliminary PCR based KoRV pol screen it was evident in the RNAseq study that they do in fact have at least partial KoRV sequences. Long read nanopore based DNA sequencing subsequently demonstrated that these sequences are a variant of the “RecKoRV” recombinant retroelements demonstrated in northern animals^49^. These are a recombination between the middle portion of an older retrotransposon in the koala genome and partial sequences of the 5’ and 3’ ends of KoRV (with the structure LTR-partial gag-central portion of PhER, - partial TM unit of env and LTR). The southern koala sequences are apparently of a different lineage to those found in the northern animals with the substitution of an unidentified piece of DNA between the KoRV and PhER sequences that is not present in the reference genome animal.

A comparison of differing sequencing methods (whole genome nanopore sequencing), the CRISPR enrichment and a PCR and nanopore sequencing strategy demonstrates that the CRISPR method produced greater read coverage and depth to resequencing the entire genome from the same animal and has the distinct advantage of being considerably cheaper (c £1000 cw £20,000). The PCR and long read sequencing in comparision was both challenging to get a PCR that worked and produced a lower read coverage and poorer homology. These sequences were also shorter than the expected 6000 Bp and likely represent mis-priming and amplification of the KoRV sequences in the PCR. This strategy also does not produce sequence information on the insertion site of the sequences. The PCR mispriming is not unexpected as the repetitive nature of the LTRs frequently results in poor PCR amplification from genomic DNA (where there are multiple copies of these ERVs) with many other studies also failing to amplify full length KoRV proviruses from koala DNA with PCR ^17,40,59^. Partial segment PCRs of the KoRV genome (LTR-gag, gag, part of *pol, env* in two parts) on DNA extracted from blood samples from SA (results presented in ^25^) demonstrated that many SA animals that test negative on the standard KoRV qPCR have at least some of the missing KoRV segments in their DNA. This indicates that there may be low copy number (likely somatic) infections of KoRV present in addition to these high copy number germline RecKoRV sequences.

Koalas with these RecKoRV variants would have been identified as KoRV negative in previous studies as the standard tests for the virus are conventional PCR or qPCR assays targeting the portion of the *pol* gene that is missing in these sequences ^4,30,51^. Other studies using KoRV *pol* PCR tests for proviral loci in DNA have also indicated that at least some southern animals have this gene but at much lower copy numbers than in QLD animals ^4^. The pattern of deletion for more ancient retroviral loci is one of loss of the *env* genes with maintenance of the *gag-pol* genes to facilitate spread within invidual cells ^60^. The replication defective variants missing their *pro-pol* genes in the current study indicate that the drivers of retroviral endogenisation in the face of an infectious virus challenge are very different to the long term ones in well adapted virus/host systems.

These RecKoRV variants are clearly replication defective and are unlikely to have colonised the genome by themselves. They may have originally arisen by being “carried” along with replication competent viruses as occurs for other retroviruses such as Rous Sarcoma Virus ^61^. It seems likely that these variants along with infectious KoRV were present before the southern animals were genetically isolated in the 1920’s and that infectious KoRV allelles either never integrated into the genome of these animals or were lost due to the genetic bottlenecks in the Southern animals^6^. The presence of the RecKoRV variants in the Victorian animals, particularly in the animal from the founder population of French Island indicates that it is likely that all southern animals have these, calling into question whether genuinely KoRV free animals exist. Examining further animals in these populations for these variants alongside genomic KoRV A is a priority. Intriguingly these insertions do not appear to be fixed between animals or populations with only a few loci shared (and none between all animals). This is comparabile to the KoRV insertion patterns seen in the northern animals ^27^ and indicates multiple colonisation events overtime. It may indicate ongoing intracellular transposition as has been hypothesised as the mechanism for the proliferation of defective variants in older endogenised retroviruses in other species ^60^. It is also possible that depth of coverage in some animals has missed some loci and follow up studies, including a larger number of animals will be essential to confirm the distribution of these defective loci across the southern koala population.

The host genetic restriction in the SA population may also have resulted in animals with viral receptor allelles that are unable to bind infectious KoRV, restricting infectious virus replication and transmission and preventing endogenisation of infectious KoRV. This situation occurs in several mouse strains resistant to certain murine leukaemia virus strains ^62^, though to date there are no known variations between southern and northern koalas for the KoRV A and B receptors, Pit1, and THTR1 and our transcriptomics screen of the two populations did not highlight these genes as varying between northern and southern animals ^6 44,47^. It is also possible that mutations in other genes important in retroviral replication (such as retroviral restriction factors) differ between the two populations resulting in restricted replication in the SA animals, although these were not obvious in our genomic screen ^6^ and this remains to be explored.

Blockade of infectious retroviruses by defective variants has been reported for several other mammalian endogenous/exogenous retroviruses. Receptor blockade by defective Env proteins occurs in Jaagsietke sheep retrovirus (JSRV)^63^, in part explaining the tissue tropism of the exogenous virus for tissues where the endogenous variants are not expressed. Endogenous JSRV loci also exert a further block on exogenous viral replication at the viral assembly stage, where defective Gag proteins from the ERV loci are packaged along with infectious variants preventing the viral particles from being packaged and transported correctly for viral release from the cell. Receptor blockade by endogenous Env proteins has also been reported in Murine Leukaemia virus variants in mice, along with a Gag mediated block at the pre-integration step of viral replication ^64^.

In this respect a number of lncRNAs were identified downstream of KoRV inserts that may play a regulatory function in expression of genes in the reverse orientation of KoRV insert sites. However the distance between each of these inserts and the associated genes is notable. One example of this is the XPR1 gene (Xenotropic and polytropic retrovirus receptor 1) which is a receptor for certain gammaretroviruses, at which two koalas (Koala 01 and Koala 03) have inserts (K01 – 48 reads; K03 – 47 reads) 500 kb upstream from the lncRNA.

While we do not yet know which of these scenarios is responsible for the marked difference in KoRV profiles between northern and southern animals, they raise the intriguing possibility that these replication defective transcripts may be interfering in some way with the full length virus variants completing their replication cycle. Future work will need to include in vivo studies of the truncated variants identified here and whether these variants do (and at what stage) blockade infectious virus replication.

It is also possible that as the southern animals (at least the ones in this study) are not born with endogenised KoRV A, they are not immune tolerised to the virus and are more able to mount an effective immune response to it. This would potentially explain the variations in antibody profiles against KoRV A evident between northern and southern animals and the very much lower KoRV induced disease prevalence between the two populations^25,65,66^.

This study does not resolve the issue of which (if any) of the identified KoRV envelope variants is the transmissible version of the virus. As has been reported in many other studies ^34,40,44,46^ our northern animals display considerable variation in their KoRV envelope variants as would be expected for a infectious replicating retrovirus. Our SA animals (with the exception of one animal), display a much more limited env variant diversity (where there are detectable reads at all) with animals expressing env genes limited to variants A, D,and E. Animal Z was the only SA animal with reads other than these three variants. We have previously reported that SA animals (whether KoRV A positive or not) display a reduced viral load and diversity compared with their QLD counterparts ^36^. It may be that these KoRV positive animals represent those with exogenous rather than endogenous KoRV as has been posited several times ^67^and are better able to control virus replication.

The discovery of these replication defective KoRV sequences in SA animals has opened up a number of intriguing implications for both controlling disease in koala populations and the drivers of retroviral endogenisation in their hosts. The hypothesis that the replication defective variants may blockade infectious KoRV replication, if substantiated, opens up the option to use selective breeding to re-introduce this trait into the KoRV susceptible northern population, though this would need to be done with caution given the presence of other deleterious genetic mutations such as those responsible for the high incidence of oxalate nephrosis ^68^ in southern animals.

## Acknowledgements

This project was funded by the Queensland Department of the Environment and Heritage Koala Research Grant Programme 2012. NS was also supported by a Keith Mackie Lucas travel scholarship from the University of Queensland. Koalas for post mortem were accessed through the Mogill Wildlife hospital (QLD Department of the Environment and Heritage Protection) and the Adelaide Koala and Wildlife Hospital, Plympton, South Australia and Fauna Rescue of South Australia Inc. AL was supported by the VESKI Victoria Fellowship 2018. Long leat Safari park has also received support through the non-profit Koala Life foundation for work with South Australian koalas.

## Supplementary Data

**Supplementary Data** 1: Details of the animals included in this study: N/A = sample not available for testing, region of origin, QLD= Queensland, SA= South Australia, sex, M= Male, F= female, tooth wear class (age classification on a 7 point scale ^69^), KoRV proviral status (pol gene PCR on DNA from whole blood), lymph node RNA quality and NGS read details: concentration, A260/280 and A260/230 ratios, RIN value, number of paired raw reads and number of trimmed reads from each sample.

**Supplementary Data 2:** Table of sequence location and naming of KoRV and RecKoRV sequences from the koala reference genome ^70^

**Supplementary Data 3:** Fast file of sequences of KoRV and RecKoRV from the koala reference genome ^70^

**Supplementary Data 4:** Estimated counts of KoRV envelope variants for individual animals, Column A = Sequence read archive (SRA) identifier,Column B = Koala ID as per Supplementary file 1, Column C, D, E: TPM for KoRV A, *gag, pro/pol* and *env* (Genbank number AF151794), Column F-M = KoRV envelope variant based on the first 575 nucleotides of the envelope variants (B-I). Column N= State of origin, Column O= KoRV pol gene PCR on whole blood DNA.

**Supplementary Data 5:** Normalised expression Log10(estimated counts) of KoRV A complete *env* gene and the 575 nucleotides of the hypervariable region of the envelope variants (B-I) for animals from tooth wear (age) class 4. Box and whisker plots show the median and interquartile ranges (box) and minimum/maximum expression (whiskers) of groups. Data for individual animals within a group are shown by circles. QLD animals in blue and SA animals in orange. Envelope variants with significantly different expression between QLD and SA groups marked with black bars (** = P<0.001, *= P<0.005)

**Supplementary Data 6:** Sequence similarity alignment generated using EasyFig ^71^. Representative assemblies from each of Koala 1, Koala 3, and Koala 4 were compared using Blast, with regions with an identity of at least 75% between sequences connected and coloured by identity value. The location in the koala genome for each of the four assemblies is denoted by the koala reference genome contig accession number in the title for each sequence. Annotated fragments of sequence regions (PhER 5’ and PhER 3’) or incomplete genes (*gag, env*) are denoted with jagged lines at the 5’ or 3’ end of the annotation. K01 SA1 NW018344210 has a deletion seen in 50% of the assembled inserts further truncating the gag gene compared to the other representative RecKoRV assemblies. This deletion ranged from ~400 – 500 bases, depending on the assembly.

