## Supplementary data 5 for "Differential and defective expression of Koala Retrovirus indicate complexity of host and virus evolution"

**Supplementary figure files:**

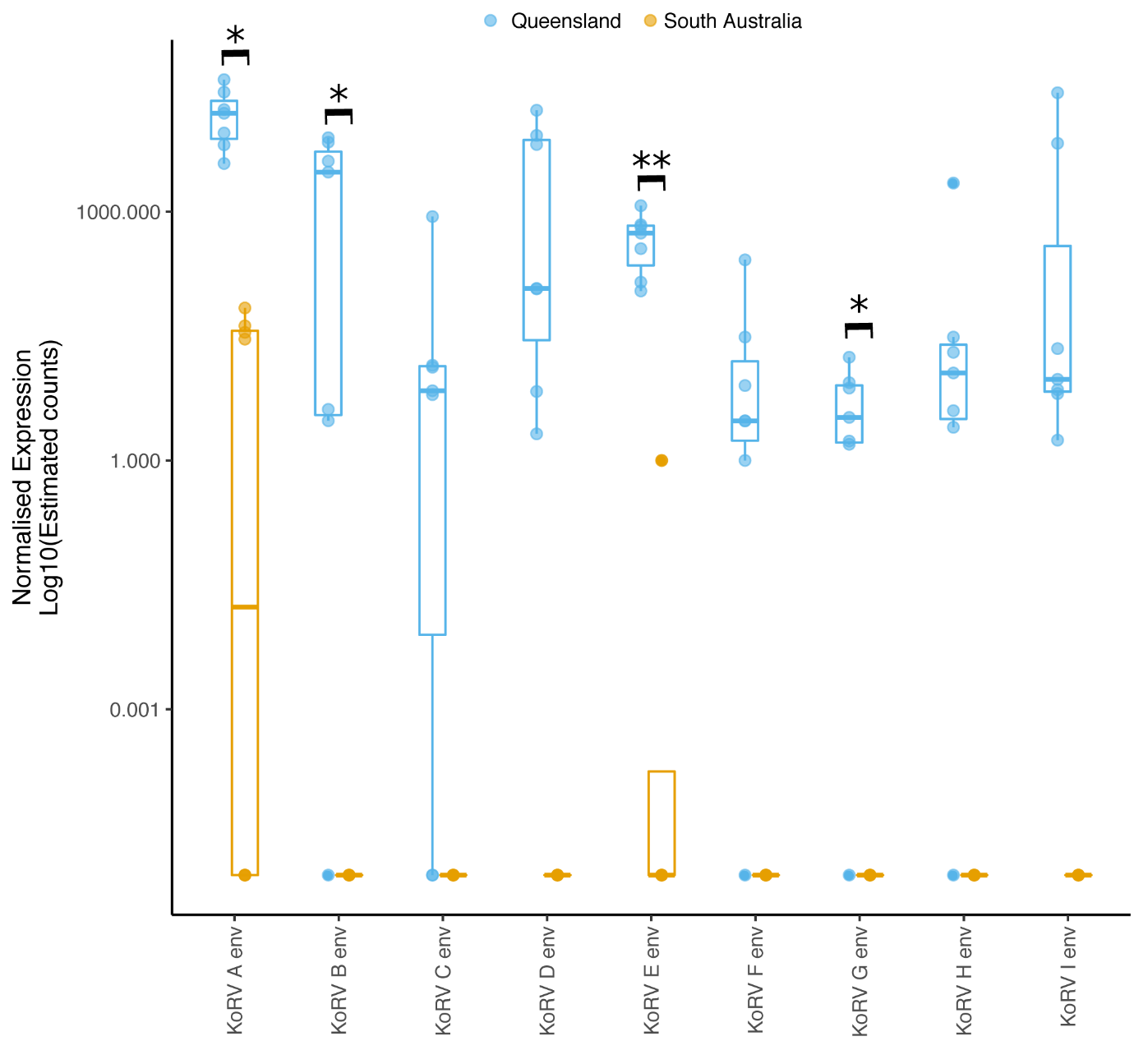

**Supplementary data 5:** Normalised expression Log10(estimated counts) of KoRV A complete *env* gene and the 575 nucleotides of the hypervariable region of the envelope variants (B-I) for animals from tooth wear (age) class 4. Box and whisker plots show the median and interquartile ranges (box) and minimum/maximum expression (whiskers) of groups. Data for individual animals within a group are shown by circles. QLD animals in blue and SA animals in orange. Envelope variants with significantly different expression between QLD and SA groups marked with black bars (** = P<0.001, *= P<0.005 )
