## Supplementary figures and images for "Differential and defective expression of Koala Retrovirus indicate complexity of host and virus evolution"

### Supplementary data 6

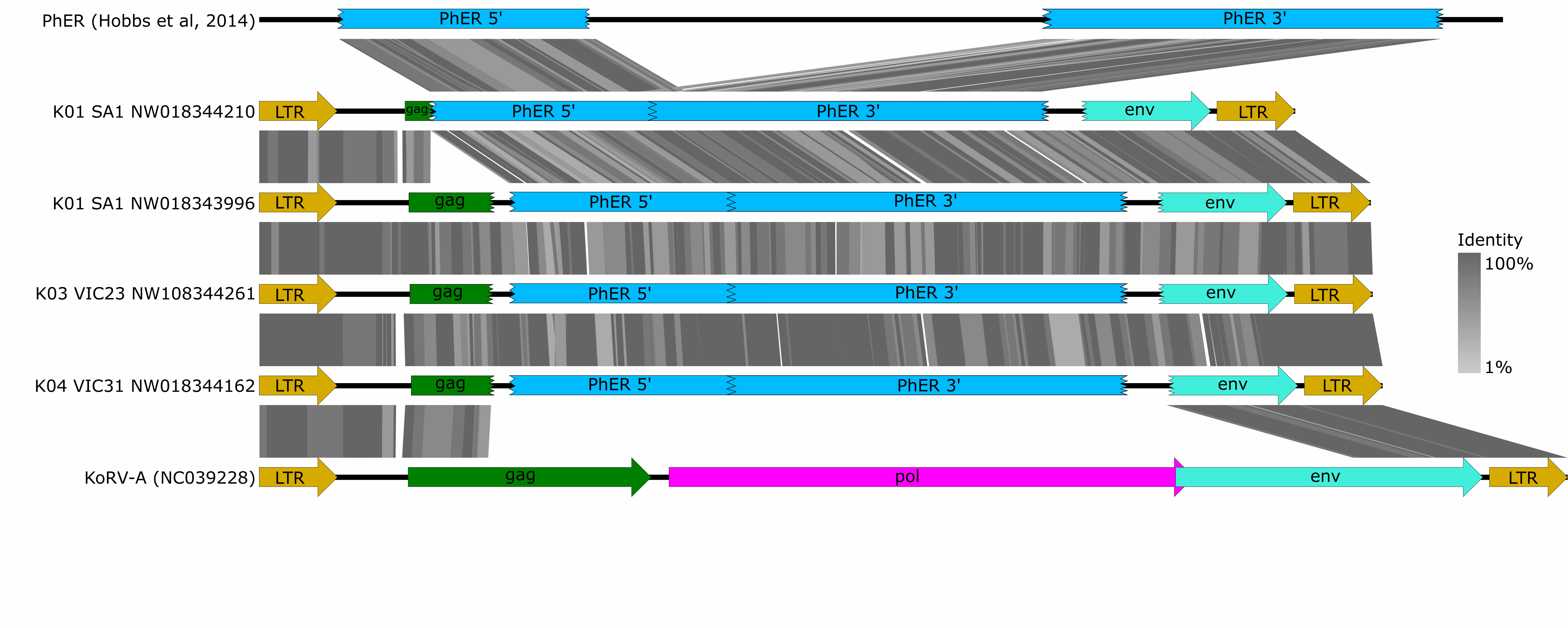
